# Aligning Model and Macaque Inferior Temporal Cortex Representations Improves Model-to-Human Behavioral Alignment and Adversarial Robustness

**DOI:** 10.1101/2022.07.01.498495

**Authors:** Joel Dapello, Kohitij Kar, Martin Schrimpf, Robert Geary, Michael Ferguson, David D. Cox, James J. DiCarlo

## Abstract

While some state-of-the-art artificial neural network systems in computer vision are strikingly accurate models of the corresponding primate visual processing, there are still many discrepancies between these models and the behavior of primates on object recognition tasks. Many current models suffer from extreme sensitivity to adversarial attacks and often do not align well with the image-by-image behavioral error patterns observed in humans. Previous research has provided strong evidence that primate object recognition behavior can be very accurately predicted by neural population activity in the inferior temporal (IT) cortex, a brain area in the late stages of the visual processing hierarchy. Therefore, here we directly test whether making the late stage representations of models more similar to that of macaque IT produces new models that exhibit more robust, primate-like behavior. We conducted chronic, large-scale multi-electrode recordings across the IT cortex in six non-human primates (rhesus macaques). We then use these data to fine-tune (end-to-end) the model “IT” representations such that they are more aligned with the biological IT representations, while preserving accuracy on object recognition tasks. We generate a cohort of models with a range of IT similarity scores validated on held-out animals across two image sets with distinct statistics. Across a battery of optimization conditions, we observed a strong correlation between the models’ IT-likeness and alignment with human behavior, as well as an increase in its adversarial robustness. We further assessed the limitations of this approach and find that the improvements in behavioral alignment and adversarial robustness generalize across different image statistics, but not to object categories outside of those covered in our IT training set. Taken together, our results demonstrate that building models that are more aligned with the primate brain leads to more robust and human-like behavior, and call for larger neural data-sets to further augment these gains.

## 1 Introduction and Related Work

Object recognition models have made incredible strides in the last ten years, [Krizhevsky et al., 2012, Szegedy et al., 2014, Simonyan and Zisserman, 2014, He et al., 2015b, Dosovitskiy et al., 2020, Liu et al., 2022] even surpassing human performance in some benchmarks [He et al., 2015a]. While some of these models bear remarkable resemblance to the primate visual system [Daniel L. Yamins, 2013, Khaligh-Razavi and Kriegeskorte, 2014, Schrimpf et al., 2018, 2020], there remain a number of important discrepancies. In particular, the output behavior of current models, while coarsely aligned with primate object confusion patterns, does not fully match primate error patterns on individual images [Rajalingham et al., 2018, Geirhos et al., 2021]. In addition, these same models can be easily fooled by adversarial attacks – targeted pixel-level perturbations intentionally designed to cause the model to produce the wrong output[Szegedy et al., 2013, Carlini and Wagner, 2016, Chen et al., 2017, Rony et al., 2018, Brendel et al., 2019], whereas primate behavior is thought to be more robust to these kinds of attacks. This is an important unsolved problem in engineering artificial intelligence systems; the deviance between model and human behavior has been studied extensively in the machine learning community, often from the perspective of safety in real-world deployment of computer vision systems [Das et al., 2017, Liu et al., 2017, Xu et al., 2017, Madry et al., 2017, Song et al., 2017, Dhillon et al., 2018, Buckman et al., 2018, Guo et al., 2018, Michaelis et al., 2019]. From a neuroscience perspective, behavioral differences like these point to different underlying mechanisms and feature representations used for object recognition between the artificial and biological systems, meaning that our scientific understanding of the mechanisms of visual behavior remains incomplete. In this work, we tested the hypothesis that we could achieve engineering/AI gains (more robustness) by seeking scientific gains (more accurate models of the brain mechanisms).

Incorporating neurophysiological constraints into models to make them behave more in line with primate visual behavior is an active field of research [Marblestone et al., 2016, Lotter et al., 2016, Nayebi and Ganguli, 2017, Guerguiev et al., 2017, Hassabis et al., 2017, Lindsay and Miller, 2018, Tang et al., 2018, Kar et al., 2019, Kubilius et al., 2019, Li et al., 2019, Hasani et al., 2019, Sinz et al., 2019, Zador, 2019, Geiger et al., 2022]. Previously, Dapello et al. [2020] demonstrated that convolutional neural network (CNN) models with early visual representations that are more functionally aligned with the early representations of primate visual processing tended to be more robust to adversarial attacks. This correlational observation was turned into a causal test, by simulating a primary visual cortex at the front of CNNs, which was indeed found to improve performance across a range of white box adversarial attacks and common image corruptions. Likewise, several recent studies have demonstrated that training models to classify images while also predicting [Safarani et al., 2021] or having similar representations [Federer et al., 2020] to early visual processing regions of primates, or even mice [Li et al., 2019], has a positive effect on generalization and robustness to adversarial attacks and common image corruptions.

Here, we extend this line of research to investigate the effects of incorporating biological knowledge of the neural representations in the IT cortex – a late stage visual processing region of the primate ventral stream, which critically supports primate visual object recognition [DiCarlo et al., 2012, Majaj et al., 2015]. To do so, we use neural recordings performed across the IT cortex of six rhesus macaque monkeys (some obtained in the study of other scientific questions [Majaj et al., 2015, Kar et al., 2019], and others recorded specifically for this work), which we divide into three training animals and three held-out testing animals for validation. We then develop a method to align the late layer “IT representations” of a base object recognition model (CORnet-S [Kubilius et al., 2019] pre-trained on ImageNet [Deng et al., 2009] and naturalistic, grey-scale “HVM” images [Majaj et al., 2015]) to the biological IT representation while the model continues to be optimized to perform classification of the dominant object in each image. We do not view these optimization procedures as having any relationship to biological learning, but simply as a way to discover models that are more brain-like than current existing models. Finally, using this strategy, we generate a suite of models under a variety of different optimization conditions and measure their IT alignment on held out animals, their alignment with human behavior, and their robustness to a range of adversarial attacks, in all cases on at least two image sets with distinct statistics as shown in figure 1.

**Figure 1:**
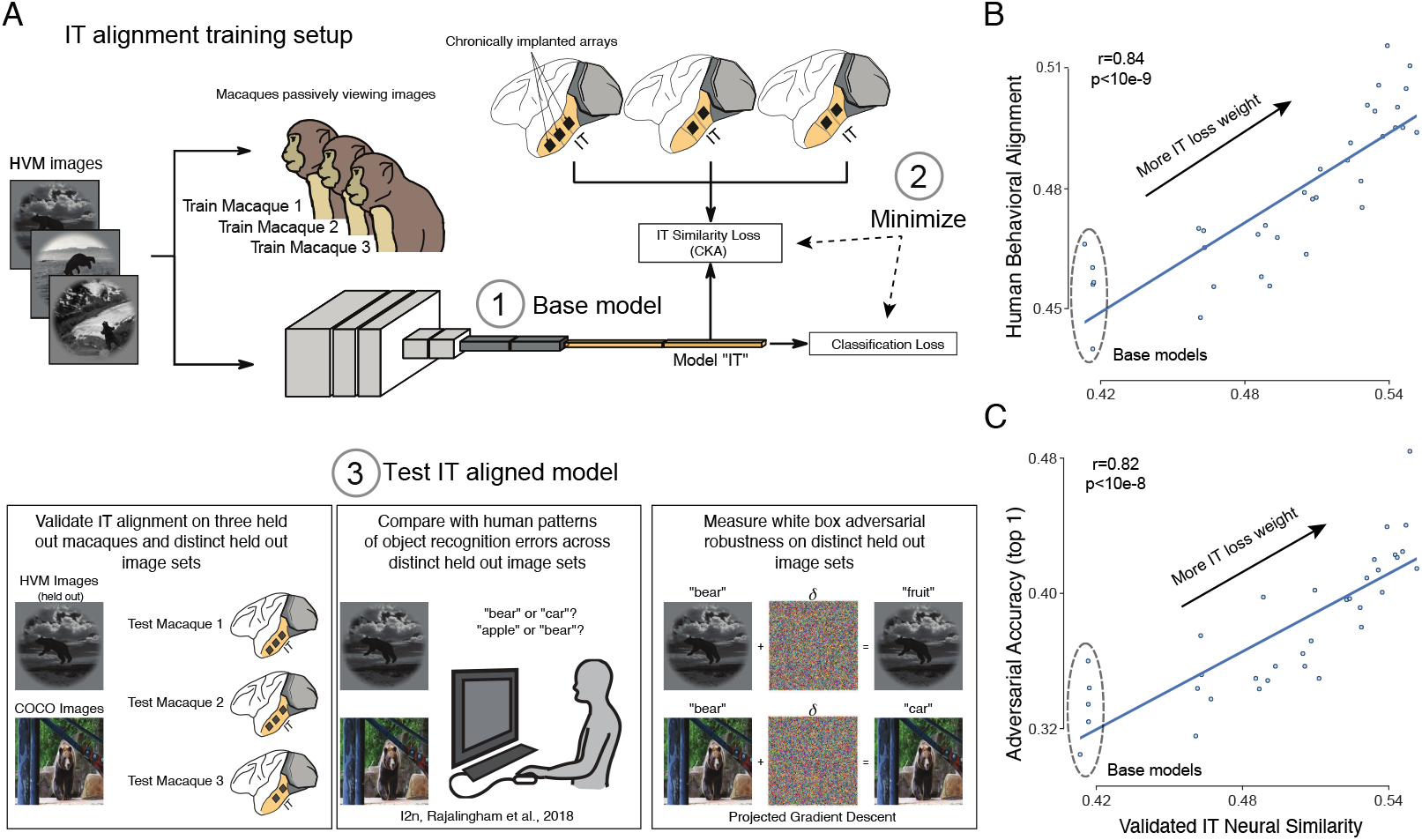
Aligning model IT representations with primate IT representations improves behavioral alignment and improves adversarial robustness. **A**) A set of naturalistic images, each containing one of eight different object classes are shown to a CNN and also to three different primate subjects with implanted multi-electrode arrays recording from the Inferior Temporal (IT) cortex. (1) A Base model (ImageNet pre-trained CORnet-S) is fine-tuned using stochastic gradient descent to (2) minimize the classification loss with respect to the ground truth object in each image while also minimizing a representational similarity loss (CKA) that encourages the model’s IT representation to be more like those measured in the (pooled) primate subjects. (3) The resultant IT aligned models are then frozen and each tested in three ways. First, model IT representations are evaluated for similarity to biological IT representation (CKA metric) using neural data obtained from new primate subjects – we refer to the split-trial reliability ceiled average across all held out macaques and both image sets as “Validated IT neural similarity”. Second, model output behavioral error patterns are assessed for alignment with human behavioral error patterns at the resolution of individual images (i2n, see Methods). Third, model behavioral output is evaluated for its robustness to white box adversarial attacks using an L_∞_ norm projected gradient descent attack. All three tests are carried out with: (i) new images within the IT-alignment training domain (held out HVM images; see Methods) and (ii) new images with novel image statistics (natural COCO images; see Methods), and those empirical results are tracked separately. **B**) We find that this IT-alignment procedure produced gains in validated IT neural similarity relative to base models on both data sets, and that these gains led to improvement in human behavioral alignment. n=30 models are shown, resulting from training at six different relative weightings of the IT neural similarity loss, each from five base models that derived from five random seeds. **C**) We also find that these same IT-alignment gains resulted in increased adversarial accuracy (PGD L_∞_, ϵ = 1/1020) on the same model set as in **B**. Base models trained only for ImageNet and HVM image classification are circled in grey.

We report three major findings:

1. Our method leads to improved IT representational similarity of models to brains even when measured on new animals and new images.
2. We find that gains in model IT-likeness lead to gains in human behavioral alignment.
3. Likewise we find that improved IT-likeness leads to increased adversarial robustness.

While probing the current limits of our IT-alignment procedure, we observed that the improvements in IT similarity, behavioral alignment, and adversarial robustness generalized to images with different image statistics than those in the IT training set (from naturalistic gray scale images to full color natural images) but only for object categories that were part of the original IT training set and not for held-out object categories.

## 2 Data and Methods

Here we describe the neural and behavioral data collection, the training and testing methods used for aligning model representations with IT representations, and the methods for assessing behavioral alignment and adversarial robustness.

### 2.1 Image sets

#### HVM image set: synthetic “naturalistic” images

High-quality images of single objects were generated using free ray-tracing software (http://www.povray.org), similar to [Majaj et al., 2015]. Each image consisted of a 2D projection of a 3D model (purchased from Dosch Design and TurboSquid) added to a random background. The ten objects chosen were bear, elephant, face, apple, car, dog, chair, plane, bird and zebra. By varying six viewing parameters, we explored three types of identity while preserving object variation, position (x and y), rotation (x, y, and z), and size. All images were achromatic with a native resolution of 256 × 256 pixels.

#### COCO image set: natural images (photographs)

Images pertaining to the 10 nouns, were download from http://cocodataset.org [Lin et al., 2014]. Each image was resized to 256 × 256 × 3 pixel size and presented within the central 8 deg.

### 2.2 Primate neural data collection and processing

We surgically implanted each monkey with a head post under aseptic conditions. We recorded neural activity using two or three micro-electrode arrays (Utah arrays; Blackrock Microsystems) implanted in IT cortex. A total of 96 electrodes were connected per array (grid arrangement, 400 um spacing, 4mm x 4mm span of each array). Array placement was guided by the sulcus pattern, which was visible during the surgery. The electrodes were accessed through a percutaneous connector that allowed simultaneous recording from all 96 electrodes from each array. All surgical and animal procedures were performed in accordance with National Institutes of Health guidelines and the Massachusetts Institute of Technology Committee on Animal Care. For information on the neural recording quality metrics per site, see supplemental section 1.1.

During each daily recording session, band-pass filtered (0.1 Hz to 10 kHz) neural activity was recorded continuously at a sampling rate of 20 kHz using Intan Recording Controllers (Intan Technologies, LLC). The majority of the data presented here were based on multiunit activity. We detected the multiunit spikes after the raw voltage data were collected. A multiunit spike event was defined as the threshold crossing when voltage (falling edge) deviated by less than three times the standard deviation of the raw voltage values. Our array placements allowed us to sample neural sites from different parts of IT, along the posterior to anterior axis. However, for all the analyses, we did not consider the specific spatial location of the site, and treated each site as a random sample from a pooled IT population. For information on the neural recording quality metrics per site, see supplemental section 1.1.

#### Behavioral state during neural data collection

All neural response data were obtained via a passive viewing task. In this task, monkeys fixated a white square dot (0.2°) for 300 ms to initiate a trial. We then presented a sequence of 5 to 10 images, each ON for 100 ms followed by a 100 ms gray blank screen. This was followed by a water reward and an inter trial interval of 500 ms, followed by the next sequence. Trials were aborted if gaze was not held within ±2° of the central fixation dot during any point. Each neural site’s response to each image was taken as the mean rate during a time window of 70-170ms following image onset, a window that has been previously chosen to align with the visually-driven latency of IT neurons and their quantitative relationship to object classification behavior as in Majaj et al. [2015].

### 2.3 Human behavioral data collection

We measured human behavior (from 88 subjects) using the online Amazon MTurk platform which enables efficient collection of large-scale psychophysical data from crowd-sourced “human intelligence tasks” (HITs). The reliability of the online MTurk platform has been validated by comparing results obtained from online and in-lab psychophysical experiments [Majaj et al., 2015, Rajalingham et al., 2015]. Each trial started with a 100 ms presentation of the sample image (one our of 1320 images). This was followed by a blank gray screen for 100 ms; followed by a choice screen with the target and distractor objects, similar to [Rajalingham et al., 2018]. The subjects indicated their choice by touching the screen or clicking the mouse over the target object. Each subjects saw an image only once. We collected the data such that, there were 80 unique subject responses per image, with varied distractor objects. Prior work has shown that human and macaque behavioral patterns are nearly identical, even at the image grain [Rajalingham et al., 2018]. For information on the human behavioral data collection, see supplemental section 1.2.

### 2.4 Aligning object recognition model representations with macaque IT representations

In order to align neural network model representations with primate IT representations while performing classification, we use a multi-loss formulation similar to that used in Li et al. (2019) and Federer et al. (2020). Starting with an ImageNet [Deng et al., 2009] pre-trained^1^ CORnet-S model [Kubilius et al., 2019], we used stochastic gradient descent (SGD) on all model weights to jointly minimize a standard categorical cross entropy loss on model predictions of ImageNet labels (maintained from model pre-training, for stability), HVM image labels, and a centered kernel alignment (CKA) based loss penalizing the “IT” layer of CORnet-S for having representations not aligned with primate IT representations of the HVM images. CORnet-S was selected because it already has a clearly defined layer committed to region IT, close to the final linear readout of the network, but otherwise our procedure is compatible with any neural network architecture. Meanwhile CKA, a measure of linear subspace alignment, was selected as a representational similarity measure. CKA has ideal properties such as invariance to isotropic scaling and orthonormal transformations which do not matter from the perspective of a linear readout, but sensitivity to arbitrary linear transformations [Kornblith et al., 2019] which could lead to differences from a linear readout as well as allow the network to hide representations useful for image classification but not present within primate IT. CKA ranges from 0, indicating completely non-overlapping subspaces, to 1, indicating completely aligned subspaces. We found that our best neural alignment results came from minimizing the neural similarity loss function log(1 − *CKA*(*X, Y*)), where 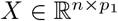 and 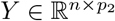 denote two column centered activation matrices with generated by showing n example images and recording p_1_ and p_2_ neurons from the IT layer of CORnet-S and macaque IT recordings respectively. The macaque neural activation matrices were generated by averaging over approximately 50 trials per image and over a 70-170 millisecond time window following image presentation. An illustration of our setup and image sets are shown in figure 1A.

### 2.5 Training and Testing Conditions

In all reported experiments, model IT representational similarity training was performed on 2880 grey-scale naturalistic HVM image representations consisting of 188 active neural sites collated from the three training set macaques for 1200 epochs. We use a batch size of 128, meaning the CKA loss computed for a random set of 128 representations for each gradient step. In order to create models with a variety of different final neural alignment scores, we add random probability 1 − *p* of dropping the IT alignment gradients and create six different sets (5 random seeds for each set) of neurally aligned models with *p* ∈ [0, 1/32, 1/16, 1/8, 1/4, 1/2, 1]. For example, the set with *p* = 0 drops all of the IT alignment gradients and thus has no improved IT alignment over the base model, while the set with *p* = 1 always includes the IT alignment gradients and similarly achieves the highest IT alignment scores (see figure 2). We also introduce a small amount of data augmentation including the physical equivalent of 0.5 degrees of jitter the vertical and horizontal position of the images, 0.5 degrees of rotational jitter, and +/-0.1 degrees of scaling jitter, assuming our model has an 8 degree field of view. The strength of these augmentations were selected to simulate natural viewing conditions.

**Figure 2:**
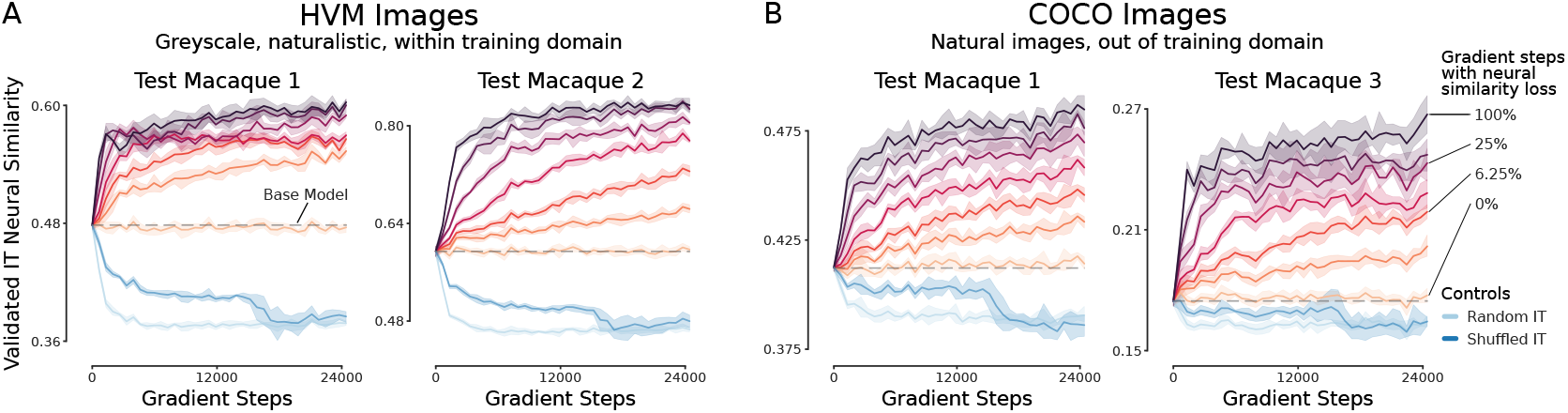
IT alignment training leads to improved IT representational similarity on held out animals and held out images across two image sets with different statistics. **A**) IT neural similarity scores (CKA, normalized by split-half trial reliability) for held out but within domain HVM images vs gradient steps is shown for two held out monkeys across seven different neural similarity loss gradient dropout rates (the darkest trace receives neural similarity loss gradients at 100% of gradient steps, while in the lightest trace neural similarity loss gradients are dropped at every step). Two control conditions are also shown: optimizing model IT toward a random Gaussian target IT matrix (random, blue) and toward an image-shuffled target IT matrix (shuffle, orange). **B**) Like **A** but for natural COCO images out of domain with respect to the training set. Grey dashed line on each plot shows the base model score for models pre-trained on ImageNet and HVM image labels with no IT representational similarity loss, which the model set with 0% of IT similarity loss gradients does not deviate significantly from. Error bars are bootstrapped confidence intervals for 5 random training seeds.

Model IT representational similarity testing was performed on a total of three held out monkeys: Monkey 1 (280 neural sites) and monkey 2 (144 neural sites) on 320 held out HVM images with statistics similar to the training distribution, and monkey 1 (237 neural sites) and monkey 3 (106 active neural sites) on 200 full color natural COCO images with different statistics than those used during training. Additional model training information can be found in supplemental section 2.

### 2.6 Behavioral Benchmarks

To characterize the behavior of the visual system, we have used an image-level behavioral metric, *i2n* [Rajalingham et al., 2018]. The behavioral metric computes a pattern of unbiased behavioral performances, using a sensitivity index: *d*′ = *Z*(*HitRate*) - *Z*(*FalseAlarmRate*), where *Z* is the inverse of the cumulative Gaussian distribution. The HitRates for i2n are the accuracies of the subjects when a specific image is shown and the choices include the target object (i.e., the object present in the image) and one other specific distractor object. So for every distractor-target pair we get a different i2n entry. A detailed description of how to compute i2n can be also found at Rajalingham et al. [2018]. The i2n behavioral benchmark was computed using the Brain-Score implementation of the i2n metric [Schrimpf et al., 2018].

### 2.7 Adversarial Attacks

For performing white box adversarial attacks, we used untargeted projected gradient descent (PGD) [Madry et al., 2017] (also referred to as the Basic Iterative Method [Kurakin et al., 2016]) with *L*_∞_, *L*_2_ norm constraints. Given an image *x*, This method uses the gradient of the loss to iteratively construct an adversarial image *x*_*adv*_ which maximizes the model loss within an *L*_*p*_ bound around *x*. Formally, in the case of an *L*_∞_ constraint, PGD iteratively computes *x*_*adv*_ as

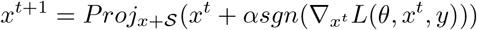

where *x* is the original image, and the Proj operator ensures the final computed adversarial image *x*_*adv*_ is constrained to the space *x* + *S*, here the *L*_∞_ ball around *x*. In the case of *L*_2_ constraints, at each iteration 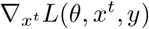 is scaled to have an *L*_2_ norm of α, and the *Proj* operator ensures the final *x*_*adv*_ is within an *L*_2_ norm ball around *x*. Unless otherwise specified, we use ∥*δ*∥_∞_ = 1/1020 constraints where δ = *x* − *x*_*adv*_ with 5 PGD iterations and a step size α = ∥*δ*∥_*p*_ /4 and report final top-1 accuracy. For more strenuous validation of our results we also compute full strength (ϵ) vs accuracy curves, and in that case we use 64 PGD steps instead of just five. The Adversarial Robustness Toolkit [Nicolae et al., 2018] was used for computing the attacks.

## 3 Results

Does aligning late stage model representations with primate IT representations lead to improvements in alignment with image-by-image patterns of human behavior or improvements in white box adversarial robustness? We start by testing if our method can generate models that are truly more IT-like by validating on held out animals and images, as this has not been previously attempted and is not guaranteed to work given the sampling limitations of neural recording experiments. We then proceed to analyze how these IT-aligned models fair on several human behavioral alignment benchmarks and a diverse set of white box adversarial attacks.

### 3.1 Direct fitting to IT neural data improves IT-likeness of models across held out animals and image sets

First, we investigated how well our IT alignment optimization procedure generalizes to IT neural similarity measurements (CKA) for two held out test monkeys on 320 held out HVM images (similar image statistics as the training set). Figure 2A shows the ceiled IT neural similarity scores for both test animals across different neural similarity loss gradient dropout rates (*p* ∈ [0, 1/32, 1/16, 1/8, 1/4, 1/2, 1]; the model marked 100% sees IT similarity loss gradients at every step, where as the model marked 0% never sees IT similarity loss gradients) as well as models optimized to classify HVM images while fitting a random Gaussian target activation matrix, or an image shuffled target activation matrix which has the same first and second order statistics as the true IT activation matrix, but scrambled image information. For both animals, we see a significant positive shift from the unfitted model (neural loss weight of 0.0), with higher relative neural loss weights generally leading to higher IT neural similarity scores. Meanwhile, both of the control conditions cause models to become less IT like, to a significant degree.

We next investigated how well our procedure generalizes from the grey-scale naturalistic HVM images to full color, natural images from COCO. Figure 2B shows the same model optimization conditions as before, but now on two unseen animal IT representations of COCO images. Like in 2A although to a lesser absolute degree, we see improvements relative to the baseline in IT neural similarity as function of the neural loss weight, and controls generally decreasing in IT neural similarity. From this, we conclude that our IT alignment procedure is able to improve IT-likeness in our models even in held out animals and across two image sets with distinct statistics.

### 3.2 Increased behavioral alignment in models that better match macaque IT

Next, we investigated how single image level classification error patterns correlate between humans and IT aligned models. To get a big picture view, we take all of the optimization conditions and validation epochs generated in figure 2A while models are training and compare IT neural similarity on the HVM test set (averaged over held out animals) with human behavioral alignment on the HVM test set. As shown in figure 3A, this analysis reveals a broad, though not linear correlation between IT neural similarity and behavioral alignment. Interestingly, we observe that the slope is at its steepest when IT neural similarity is at the highest values, suggesting that if an even higher degree of IT-alignment might result in greater increases in behavioral alignment. We also investigated whether these trends persist when we exclude the optimization on object labels from the HVM images and only optimize for IT neural similarity. To do so, we train the models on all previous conditions but without the HVM object-label loss. As shown in 3, the overall shape of the trend remains quite similar, though the absolute behavioral alignment shifts downward, indicating that the label information during training helps on the behavioral task, but is not a requirement for the trends to hold.

**Figure 3:**
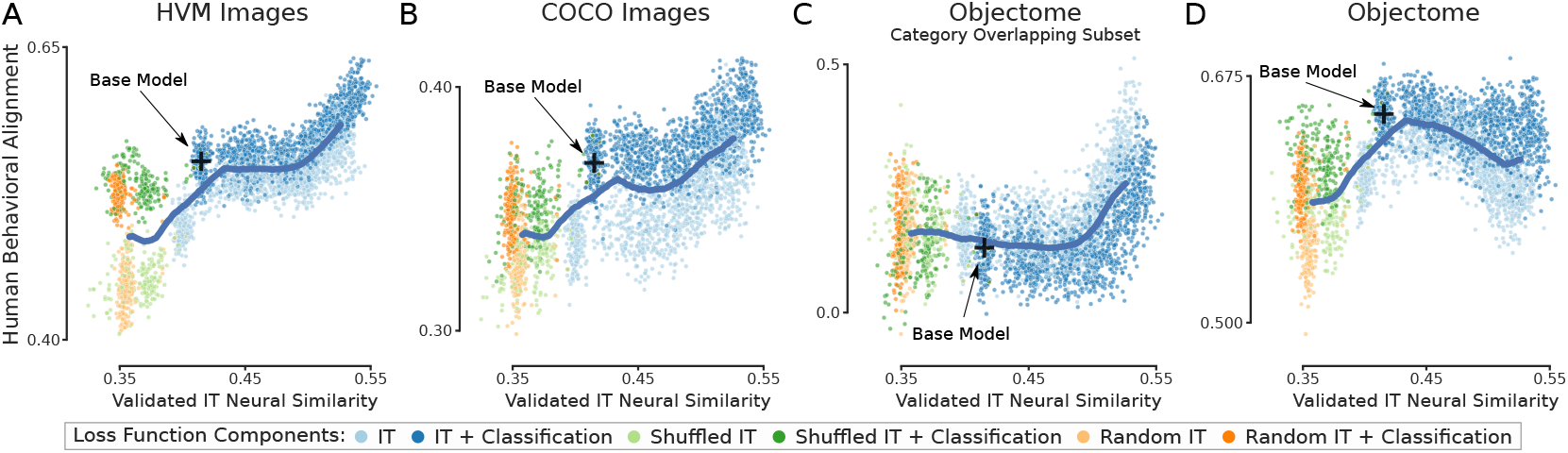
IT neural similarity correlates with behavioral alignment across a variety of optimization conditions and unseen image statistics but not on unseen object categories. **A**) Held out animal and image IT neural similarity is plotted against human behavioral alignment on the HVM image set at every validation epoch for all neural loss weight conditions, random IT target matrix conditions, and image shuffled IT target matrix conditions, in each case with or with and with out image classification loss. **B**) and **C**) Like in **A** but for the COCO image set and the Objectome image set Rajalingham et al. [2018] filtered to overlapping categories with the IT training set. **D**) The behavioral alignment for the full Objectome image set with 20 categories not covered in the IT training is not improved by the IT-alignment procedure and data used here. In all plots, the black cross represents the average base model position, and the heavy blue line is a sliding X, Y average of all conditions merely to visually highlight trends. Five seeds for each condition are plotted.

In figure 3B, we perform the same set of measurements but now focusing on the COCO image set. Consistent with the observation on COCO IT neural similarity, the behavioral alignment trend transfers to the COCO image set although the absolute magnitude of the improvements are less.

Finally, using the Brain-Score platform [Schrimpf et al., 2018], we benchmark our models against publicly available human behavioral data from the Objectome image set Rajalingham et al. [2018] which has similar image statistics to our HVM IT fitting set (with a total of 24 object categories, only four of which overlap with the training set). As demonstrated in figure 4C, when the Objectome data are filtered down to just the four overlapping categories, our most IT similar models are again the most behaviorally aligned, well above the unfit baseline and control conditions, which remain close to the floor for much of the plot. However, As shown in figure 3D, when considering all 24 object categories in the Objectome dataset, we see that the trend of increasing human behavioral alignment does not hold and our models actually begin to fair worse in terms of human behavioral alignment at higher levels of IT neural similarity. As shown in figure supp 3.1, using a linear probe to assess image class information content (measured by classification accuracy on held out representations) reveals that these models are losing class information content for the Objectome image set, which drives the decrease in behavioral alignment, as the model makes more mistakes overall than a human. Similarly, a linear probe analysis reveals minimal loss in class information in the overlapping categories. Thus, we observe that while our method leads to increased human behavioral alignment across different image statistics, it does not currently lead to improved alignment on unseen object categories.

**Figure 4:**
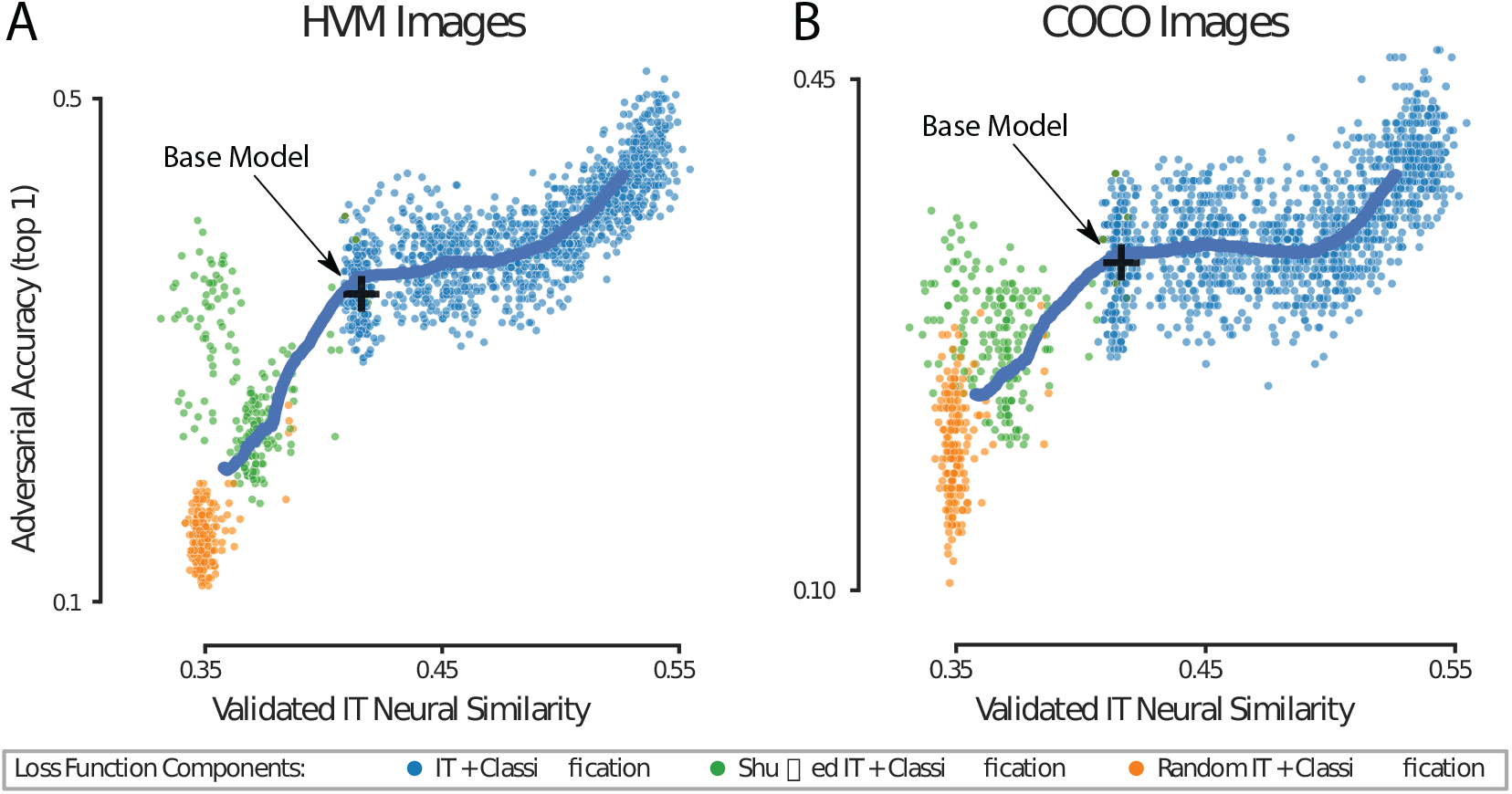
IT neural similarity correlates with improved white box adversarial robustness. **A**) held out animal and image IT neural similarity is plotted against white box adversarial accuracy (PGD *L*_∞_ *ϵ* = 1/1020) on the HVM image set measured across multiple training time points for all neural loss ratio conditions, random IT target matrix conditions, and image shuffled IT target matrix conditions. **B**) Like in **A** but for COCO images. In all plots, the black cross represents the average base model position, and the heavy blue line is a sliding X, Y average of all conditions merely to visually highlight trends. Five seeds for each condition are plotted.

### 3.3 Increased adversarial robustness in models that better match macaque IT

Finally, we evaluate our models on an array of white box adversarial attacks, to assess if models with higher IT neural similarity scores also have increased adversarial robustness. Like before, we start with a big picture analysis where we consider every evaluation epoch for all optimization conditions considered in figure 2. Again, as demonstrated in figures 4A and 4B, for both HVM images and COCO images, there is a broad though not entirely linear correlation between IT neural similarity and adversarial robustness to PGD *L*_∞_ *ϵ* = 1/1020 attacks. Like in the analysis of behavioral alignment, we also see a higher slope on the right side of the plots, where IT neural similarity is the highest, suggesting further improvements could be had if models were pushed to be more IT aligned.

In order to get a better sense of the gains in robustness, we measured the adversarial strength accuracy curves for models only trained with HVM image labels, models trained with HVM image labels and IT neural representations, and models adversarially trained on HVM labels (PGD *L*_∞_, *ϵ* = 4/255). Figure 5A shows that on held-out HVM images, IT aligned models have increased accuracy across a range of *ϵ* values for both *L*_∞_ and *L*_2_ norms, though less so than models with explicit adversarial training. However, as shown in figure 5 the same analysis on COCO images demonstrates that adversarial robustness in the IT aligned networks generalizes significantly better on unseen image statistics than the adversarially trained models, which lose clean accuracy on COCO images.

**Figure 5:**
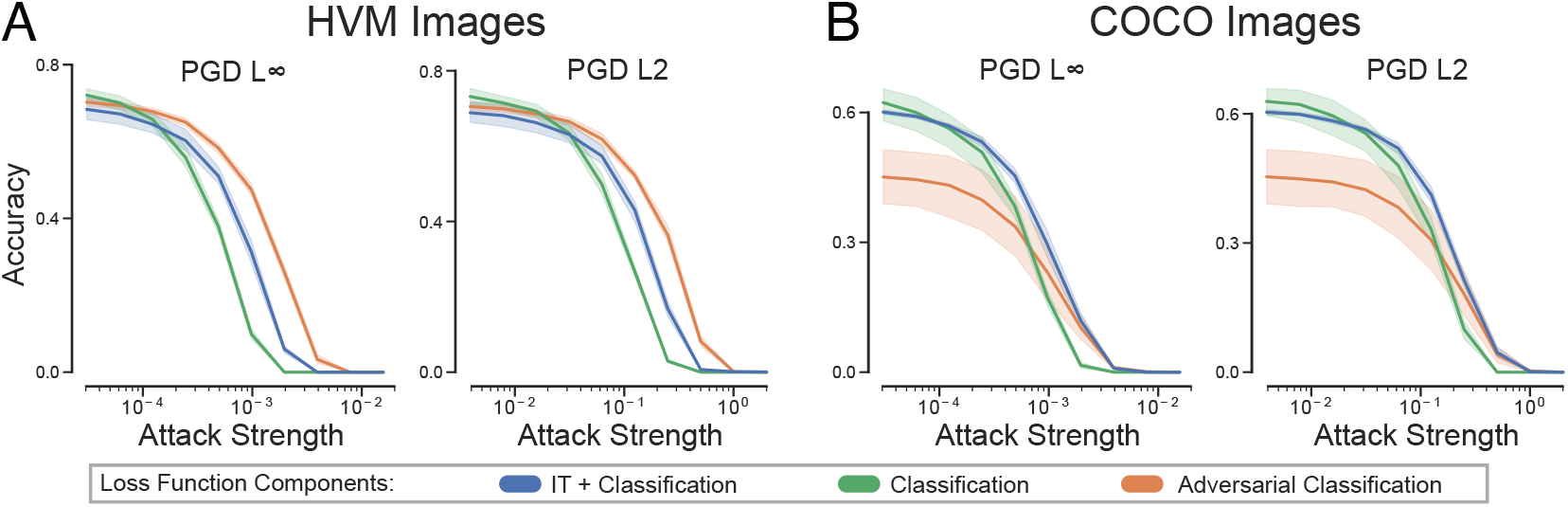
IT aligned models are more robust than standard models in and out of domain, and more robust than adversarially trained models in out of domain conditions. **A**) PGD *L*_∞_ and *L*_2_ strength accuracy curves on HVM images for standard trained networks (green) IT aligned networks (blue) and networks adversarially trained (PGD *L*_∞_ *ϵ* = 4/255) on the IT fitting image labels (orange). **B**) Like in **A** but for COCO images. Error shading represents bootstrapped 95% confidence intervals over five training seeds.

Finally, we tested the IT neural similarity of our HVM image adversarially trained models and find that they do not follow the general correlation shown in 4 for IT aligned models vs adversarial accuracy. Interestingly, the adversarially trained models are slightly more similar to IT than standard models, but significantly higher than standard models on HVM adversarial accuracy and significantly lower on COCO adversarial accuracy. We take this to indicate that there are multiple possible ways to become robust to adversarial attacks, and that adversarial training does not in general induce the same representations as IT alignment.

## 4 Discussion

Building on prior research in constraining visual object recognition models with early stage visual representations [Li et al., 2019, Dapello et al., 2020, Federer et al., 2020, Safarani et al., 2021], we report here that it is possible to better align the late stage “IT representations” of an object recognition model with the corresponding primate IT representations, and that this improved IT alignment leads to increased human level behavioral alignment and increased adversarial robustness. In particular, the results show that 1) the method used here is able to improve IT alignment in object recognition models even on held out animals and image statistics not seen by the model during the IT neural alignment training procedure, 2) models that are more aligned with macaque IT also have better alignment with human behavioral error patterns across unseen (not shown during training) image statistics but not for unseen object categories, and 3) models more aligned with macaque IT are more robust to adversarial attacks even on unseen image statistics. Interestingly however, we observed that being more adversarially robust does not lead to significantly more IT neural similarity.

These empirical observations raise a number of important questions for future research. First, we find it intriguing that aligning IT representations in our models to empirically measured macaque IT responses has no effect or even a negative effect on behavioral alignment for objects not present in the IT fitting image-set, a noteworthy limitation in our approach. We speculate that this is due to the small range of categories covered in our IT training set, which limits the span of neural representational space that those experiments were able to sample. In that regard, it would be informative to get a sense of the scaling laws [Kaplan et al., 2020] for how much neural data (in terms of images, neurons, trials, or object categories) needs to be absorbed into a model before it behaves in a truly general more human like fashion for any instance of image categories or statistics. With regard to adversarial robustness, a number of questions remain as well. While there are clear gains in robustness from our procedure, the overall magnitude is relatively small. How much adversarial robustness could we expect to gain, if we perfectly fit IT? And how much could be gained from fitting other regions of the ventral stream more precisely? These question hinges on how adversarially robust primate behavior really is, an active area of research [Guo et al., 2022, Elsayed et al., 2018, Yuan et al., 2020]. We also note that our current study do not exhaustively test the influence of various model architectures, training data, and learning algorithms (or other optimization goals) on the ability of the model to better absorb macaque IT responses during training. Finally, the question remains as to why primate visual representations are more robust than representations developed in models with the standard classification loss.

Overall, our results provide further evidence in support of the framework encouraging direct optmization of models with empiricial data from the primate brain to make them more robust and well aligned with human behavior [Sinz et al., 2019]. We also highlight that there are many compelling ways forward. In hopes that more neurophysiological constraints will further limit the model solution space to resemble what is used by the primate brain, a clear next step is to combine our IT alignment method with prior methods developed to increase alignment with early visual representations, either with a fixed V1 front end [Dapello et al., 2020], or in a more data driven manner [Safarani et al., 2021]. Beyond this, our method could be adapted to constrain recurrent models [Kubilius et al., 2019, Nayebi et al., 2018] with the recorded temporal dynamics of IT neural responses [Kar et al., 2019]. Finally, we could also test if adding neural stochasticity offers improvements in robustness here as it did in Dapello et al. [2020, 2021].

## Acknowledgments and Disclosure of Funding

We thank Sachi Sanghavi for the assistance with primate neural recording data and Kathleen Leeper for paper editing, support, and insightful discussions. This work was supported by the Semiconductor Research Corporation (SRC) and DARPA (J.D., M.S., J.J.D.), the Massachusetts Institute of Technology Shoemaker Fellowship (M.S.), Office of Naval Research grant MURI-114407 (J.J.D.), the Simons Foundation grant SCGB-542965 (J.J.D.), and Canada Research Chair Program (K.K.).

## Checklist

1. For all authors…
  a. Do the main claims made in the abstract and introduction accurately reflect the paper’s contributions and scope? [Yes] All claims made in the abstract and intro are directly supported by the figures in the paper.
  b. Did you describe the limitations of your work? [Yes] The middle paragraph of the discussion focuses on limitations of the work
  c. Did you discuss any potential negative societal impacts of your work? [No] We do not foresee any negative societal implications in IT aligning object recognition models.
  d. Have you read the ethics review guidelines and ensured that your paper conforms to them? [Yes]
2. If you are including theoretical results…
  a. Did you state the full set of assumptions of all theoretical results? [N/A] We do not use any proofs.
  b. Did you include complete proofs of all theoretical results? [N/A] We do not use any proofs.
3. If you ran experiments…
  a. Did you include the code, data, and instructions needed to reproduce the main experimental results (either in the supplemental material or as a URL)? [Yes] We include instructions to reproduce our work and will release a github repository with all code and data needed to reproduce our results upon paper acceptance.
  b. Did you specify all the training details (e.g., data splits, hyperparameters, how they were chosen)? [Yes] In the materials and methods section as well as further in the supplemental materials.
  c. Did you report error bars (e.g., with respect to the random seed after running experiments multiple times)? [Yes] All experiments were run with 5 random seeds per condition and plotted with error bars.
  d. Did you include the total amount of compute and the type of resources used (e.g., type of GPUs, internal cluster, or cloud provider)? [Yes] In supplemental material section 2, where we discuss additional model training details.
4. If you are using existing assets (e.g., code, data, models) or curating/releasing new assets…
  a. If your work uses existing assets, did you cite the creators? [Yes] We cite Brain-Score, ART, Kubilius et al. [2019], Majaj et al. [2015], and Rajalingham et al. [2018], whose code and data we use in our work.
  b. Did you mention the license of the assets? [Yes] We include code and asset licensing information where appropriate in supplemental section 4.
  c. Did you include any new assets either in the supplemental material or as a URL? [Yes] We will release new assets upon paper acceptance.
  d. Did you discuss whether and how consent was obtained from people whose data you’re using/curating? [N/A] All non-original data was taken from public sources (like published papers) where consent was not necessary
  e. Did you discuss whether the data you are using/curating contains personally identifiable information or offensive content? [N/A] Our data contains nothing of a personally identifiable nature.
5. If you used crowdsourcing or conducted research with human subjects…
  a. Did you include the full text of instructions given to participants and screenshots, if applicable? [Yes] We include relevant information in supplemental section 1.2
  b. Did you describe any potential participant risks, with links to Institutional Review Board (IRB) approvals, if applicable? [N/A] There were no risks to our participants.
  c. Did you include the estimated hourly wage paid to participants and the total amount spent on participant compensation? [Yes] We include relevant information in supplemental section 1.2

## Supplementary Material

### A Data Collection and Processing

#### A.1 Neural Data Collection and Processing

##### A.1.1 Surgical implant of chronic micro-electrode arrays

In this study we have recorded chronic, in-vivo neural responses across both left and right hemispheres of 6 specific monkeys (3 used in model training and 3 used in model testing). First, we surgically implanted each monkey with a head post under aseptic conditions. After behavioral training, we recorded neural activity using multiple 10 × 10 micro-electrode arrays (Utah arrays; Blackrock Microsystems). A total of 96 electrodes were connected per array. Each electrode was 1.5 mm long and the distance between adjacent electrodes was 400 um. Before recording, we implanted each monkey multiple Utah arrays in the IT cortex. Array placement was guided by the sulcus pattern, which was visible during surgery. The electrodes were accessed through a percutaneous connector that allowed simultaneous recording from all 96 electrodes from each array. All behavioral training and testing was performed using standard operant conditioning (water reward), head stabilization, and real-time video eye tracking. All surgical and animal procedures were performed in accordance with National Institutes of Health guidelines and the Massachusetts Institute of Technology Committee on Animal Care.

##### A.1.2 Electrophysiological Recording

During each recording session, band-pass filtered (0.1 Hz to 10 kHz) neural activity was recorded continuously at a sampling rate of 20 kHz using Intan Recording Controller (Intan Technologies, LLC). The majority of the data presented here were based on multiunit activity. We detected the multiunit spikes after the raw data was collected. A multiunit spike event was defined as the threshold crossing when voltage (falling edge) deviated by less than three times the standard deviation of the raw voltage values. Our array placements allowed us to sample neural sites from different parts of IT, along the posterior to anterior axis. However, for all the analyses, we did not consider the specific spatial location of the site, and treated each site as a random sample from a pooled IT population.

###### Monkeys used for model training

We recorded neural data from three monkeys for training the CNNs. Chronic multi-electrode recordings from these monkeys’ ventral visual cortex (recorded for various other research objectives) has been previously published [Majaj et al., 2015] [Hong et al., 2016] [Bashivan et al., 2019] [Rajalingham et al., 2020].

###### Monkeys used for model testing

We recorded neural data from three monkeys for testing the jointly optimized models. Chronic multi-electrode recordings from these monkeys’ ventral visual cortex (recorded for various other research objectives) has been previously published [Kar et al., 2019] [Kar and DiCarlo, 2021] [Bashivan et al., 2019] [Rajalingham et al., 2020]

##### A.1.3 Neural recording quality metrics per site

Visual drive per neuron 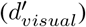: We estimated the overall visual drive for each electrode. This metric was estimated by comparing the selected image responses of each site to a blank (gray screen) response.

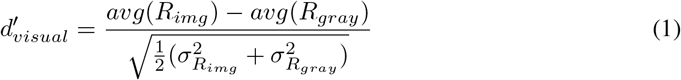

Image rank-order response reliability per neural site 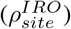: To estimate the reliability of the responses per site, we computed a spearman-brown corrected, split half (trial-based) correlation between the rank order of the image responses (all images).

Selectivity per neural site: For each site, we measured selectivity as the d’ for separating that site’s best (highest response-driving) stimulus from its worst (lowest response-driving) stimulus. d′ was computed by comparing the response mean of the site over all trials on the best stimulus as compared to the response mean of the site over all trials on the worst stimulus, and normalized by the square-root of the mean of the variances of the sites on the two stimuli:

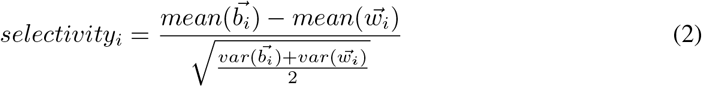

Where 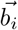 is the vector of responses of site i to its best stimulus over all trials and 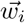 is the vector of responses of site i to its worst stimulus. We computed this number in a cross-validated fashion, picking the best and worst stimulus on a subset of trials and then computing the selectivity measure on a separate set of trials, and averaging the selectivity value of 50 trial splits.

Inclusion criterion for neural sites: For our analyses, we only included the neural recording sites that had an overall significant visual drive 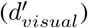, an image rank order response reliability 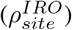 that was greater than 0.6 and a selectivity score that was greater than 1. Given that most of our neural metrics are corrected by the estimated noise at each neural site, the criterion for selection of neural sites is not that critical, and it was mostly done to reduce computation time by eliminating noisy recordings.

###### A.2 Human Data Collection

We measured human behavior (from 88 subjects) using the online Amazon MTurk platform which enables efficient collection of large-scale psychophysical data from crowd-sourced “human intelligence tasks” (HITs). The reliability of the online MTurk platform has been validated by comparing results obtained from online and in-lab psychophysical experiments [Majaj et al., 2015] [Rajalingham et al., 2015]. Each trial started with a 100 ms presentation of the sample image (that contained a visual object embedded in a scene). This was followed by a blank gray screen for 100 ms; followed by a choice screen with the target and distractor objects (similar to [Rajalingham et al., 2018]). The subjects indicated their choice by touching the screen (for touch screen tablets) or clicking the mouse over the target object (for desktop computers). Each subjects saw an image only once. We collected the data such that, there were 80 unique subject responses per image, with varied distractor objects.

Human participants were compensated at a rate of approximately 4 USD per hour. The total amount spent was approximately 300 USD.

Human participants were greeted with the following messages upon clicking on the task on Amazon Mechanical Turk.

###### Welcome page instructions

“This HIT is part of a MIT scientific research project. Your decision to complete this HIT is voluntary. There is no way for us to identify you. The only information we will have, in addition to your responses, is the time at which you completed the survey. The results of the research may be presented at scientific meetings or published in scientific journals. Clicking on the ‘SUBMIT’ button on the bottom of this page indicates that you are at least 18 years of age and agree to complete this HIT voluntarily.

NOTE: Please close all other programs/taps while running this task to get the optimal system performance. Users on a suboptimal system can experience glitches that will lead to rejection. Also, low scores on this task will lead to rejection: make sure to read this instruction! Thank you for your interest! You are contributing to ongoing vision research at the Massachusetts Institute of Technology and the McGovern Institute for Brain Research. This task will require you to look at images on your computer screen and click to indicate a response for up to about 15 minutes, although we expect this to take about 5-10 minutes. If you cannot meet these requirements for any reason, or if doing so could cause discomfort or injury to you, do not accept this HIT. We encourage you to try a little bit of this HIT before accepting to ensure it is compatible with your system. If you think the task is working improperly, your computer may be incompatible. We recommend this task for those who are interested in contributing to scientific endeavors. Your answers will help MIT researchers better understand how the brain processes visual information.”

###### Task instructions

“Please fixate at the center ‘+’ sign which will appear once you press any key. Then, an image will be shown in the center of the screen. The image will contain an object. This will be followed by a Choice Screen containing two objects. Please choose the option [Click on the object] that you think was present in the image”

### B Additional Model Training and Testing Details

In order to align model representations with primate IT representations while performing a classification task, we start by loading an ImageNet [Deng et al., 2009] pre-trained CORnet-S [Kubilius et al., 2019] as a base model. We add 8 new classification logits for the HVM image classes, and model hooks to extract representations from the first time step of the “IT” layer of CORnet-S. Next, we resume ImageNet training of the CORnet-S using stochastic gradient descent with a batch size of 128, a learning rate of 0.001, 0.9 momentum, and weight decay of 0.0001, but now for every gradient step we also include a batch of HVM images. For each HVM batch, a cross entropy loss is applied to the 8 new logits for classifying the HVM images, and a *log*(1 − *CKA*(*X, Y*)) loss is applied, where *X* is the activation matrix for the batch of representations extracted from CORnet-S “IT” layer and Y is the activation matrix of the corresponding image representations recorded from IT in the three training macaques. To generate a diversity of IT similarity in our models, we introduce random dropout of the CKA similarity loss which we call a neural loss ratio. A neural loss ratio of 1:1 corresponds to all HVM images having both a neural similarity loss and an HVM classification loss, while a neural loss ratio of 0.5:1 means that 50% of the time, the HVM batch only includes the HVM classification loss and no neural similarity loss. A neural loss ratio of 0:1 means that the model never has neural similarity gradients, only HVM classification gradients during the HVM batches. Finally, for experiments like 3 where we explore the effect of HVM label information on behavioral alignment, the HVM classification loss is set to 0 on all HVM batches. At no point during the training procedure does the model see COCO images or IT representations of COCO images.

In all reported experiments, model IT representational similarity training was performed on 2880 grey-scale naturalistic HVM image representations consisting of 188 active neural sites collated from the three training set macaques for 1200 epochs, or 26400 gradient steps on QUADRO RTX6000 GPUs, which took approximately 5 hours per training run. We introduce a small amount of data augmentation including the physical equivalent of 0.5 degrees of jitter the vertical and horizontal position of the image, 0.5 degrees of rotational jitter, and +/-0.1 degrees of scaling jitter, selected to simulate natural viewing conditions. All models are in pytorch [Paszke et al., 2017], and training and testing is performed using pytorch lightning [Falcon et al., 2019].

Model IT representational similarity testing was performed on a total of three held out monkeys: Monkey 1 (280 neural sites) and monkey 2 (144 neural sites) on 320 held out HVM images with statistics similar to the training distribution, and monkey 1 (237 neural sites) and monkey 3 (106 active neural sites) on 200 full color natural COCO [Lin et al., 2014] images with different statistics than those used during training. IT neural similarity was computed as 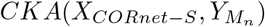 where *X*_*CORnet*−*S*_ denotes the activation matrix of CORnet-S time step zero responses from the “IT” layer and 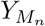 is the activation matrix o−f test macaque n responses to all test images from an image set (HVM or COCO). The CKA score is computed for each test animal individually and normalized by the split half trial reliability of the animal’s recordings, ie *CKA*(*X, Y*) where *X* is the time and trial averaged recordings for 50% of the images presentations and Y is the time and trial averaged recordings for the other 50% of image presentations, bootstrapped over 10 random splitting seeds. Finally, the we average the individually computed and ceiled scores of each animal to give the Validated IT Neural Similarity scores reported in all experiments.

### C Additional Experimental Results

Here, to investigate why the behavioral alignment on Objectome Rajalingham et al. [2018] images decreases for higher levels of validated IT neural similarity, we use a linear probe to assess the (linear) class information content of Objectome image representations. Briefly, like for computing i2n behavioral alignment, feature representations are extracted for all image from the final average pooling layer of the network and partitioned into a train and test set. A linear classifier is fit to the train partition representations, and the classification error on the test set is reported. Results are shown in figure A.1. As can be seen, in the case of the full Objectome image set, classification accuracy decreases for higher values of IT neural similarity. Meanwhile, in the category overlapping subset, there is minimal loss in overall classification accuracy. Since i2n behavioral alignment is implicitly reliant on raw classification accuracy, this is at least one source of the decreasing behavioral alignment for Objectome images. Still, it is surprising to us that the network representations for categories beyond the IT fitting set are being degraded, and more research is needed to understand exactly what is happening to the underlying representations.

**Figure A.1:**
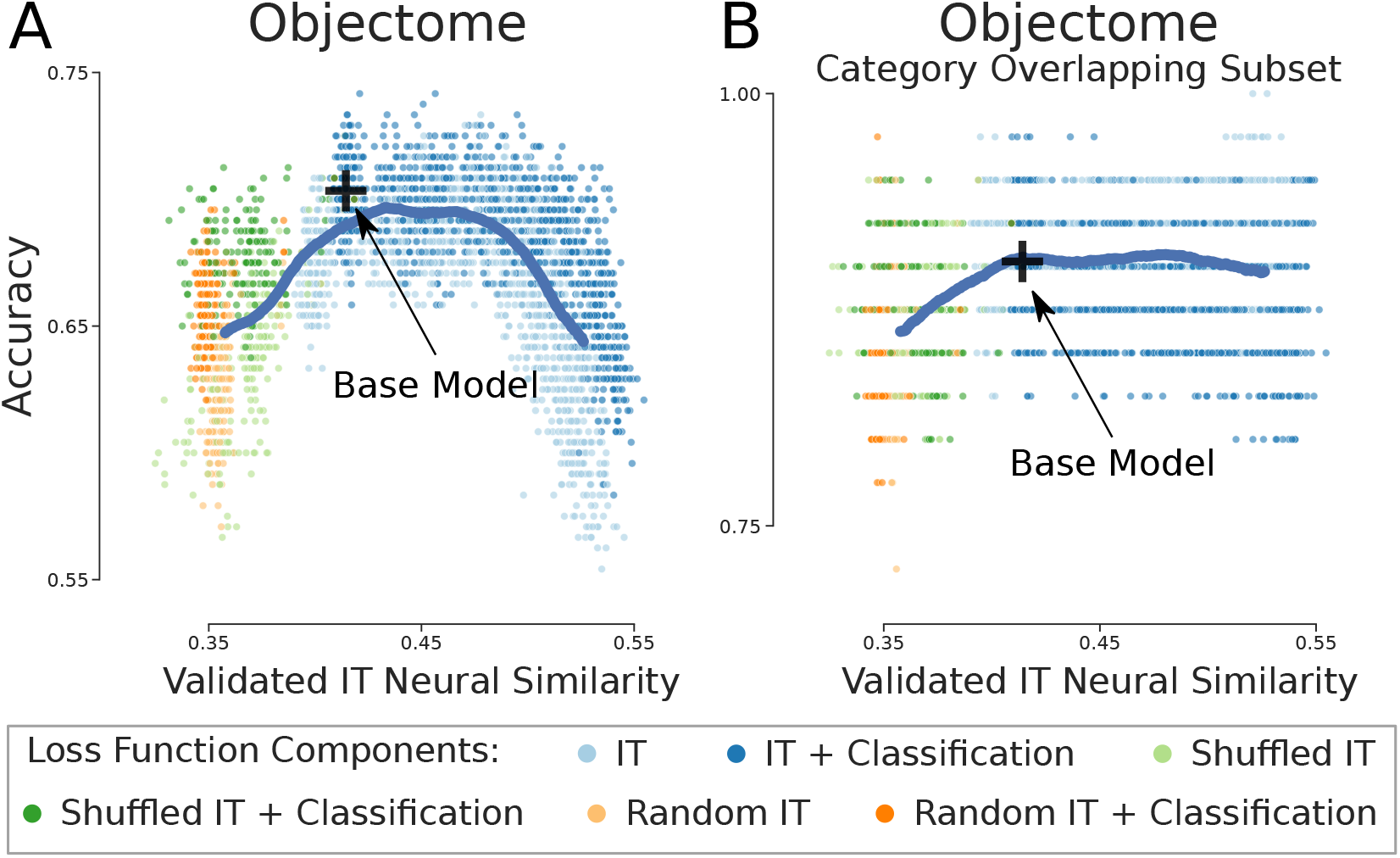
Higher levels of validated IT neural similarity are associated with decreasing classification accuracy on Objectome images over all, but not Objectome images with categories overlapping the IT training set categories. **A**) Classification accuracy versus validated IT neural similarity for all model optimization conditions at all validation checkpoints reveals decreasing accuracy on the full Objectome image set which contains 20 categories not overlapping with the IT training set. **B**) Like in **A** but for the Objectome image set filtered to overlapping categories with the IT training set has relatively stable classification accuracy. In all plots, the black cross represents the average base model position, and the heavy blue line is a sliding X, Y average of all conditions merely to visually highlight trends. Five seeds for each condition are plotted.

### D Licensing Information

The MS COCO images dataset [Lin et al., 2014] is licensed under a Creative Commons Attribution 4.0 License. Adversarial Robustness Toolbox [Nicolae et al., 2018], Brain-Score [Schrimpf et al., 2018], Brain-Score associated datasets are under the MIT License. Pytorch [Paszke et al., 2017] is under a BSD license. Pytorch Lightning [Falcon et al., 2019] is under an Apache 2.0 license. CORnet-S [Kubilius et al., 2019] is under a GNU General Public License. ImageNet [Deng et al., 2009] is under a BSD 3-Clause License.

## Footnotes

1 the pre-trained version was selected as a starting point because of the relatively small number of training samples in our dataset [Riedel, 2022].

## Notes

### Competing Interest Statement

The authors have declared no competing interest.

## References

P. Bashivan, K. Kar, and J. J. DiCarlo. Neural population control via deep image synthesis. Science, 364(6439), May 2019.

W. Brendel, J. Rauber, M. Kümmerer, I. Ustyuzhaninov, and M. Bethge. Accurate, reliable and fast robustness evaluation. July 2019.

J. Buckman, A. Roy, C. Raffel, and I. Goodfellow. Thermometer encoding: One hot way to resist adversarial examples. Feb. 2018.

N. Carlini and D. Wagner. Towards evaluating the robustness of neural networks. Aug. 2016.

P.-Y. Chen, Y. Sharma, H. Zhang, J. Yi, and C.-J. Hsieh. EAD: Elastic-Net attacks to deep neural networks via adversarial examples. Sept. 2017.

C. C. J. J. D. Daniel L. Yamins, Ha Hong. Hierarchical modular optimization of convolutional networks achieves representations similar to macaque IT and human ventral stream. Advances in Neural Information Processing Systems 26 (NIPS 2013), 2013.

J. Dapello, T. Marques, M. Schrimpf, F. Geiger, D. D. Cox, and J. J. DiCarlo. Simulating a primary visual cortex at the front of CNNs improves robustness to image perturbations. page Advances in Neural Information Processing Systems 33 (NeurIPS 2020). Neurips, June 2020.

J. Dapello, J. Feather, H. Le, T. Marques, D. Cox, J. McDermott, J. J. DiCarlo, and S. Chung. Neural population geometry reveals the role of stochasticity in robust perception. Adv. Neural Inf. Process. Syst., 34, 2021.

N. Das, M. Shanbhogue, S.-T. Chen, F. Hohman, L. Chen, M. E. Kounavis, and D. H. Chau. Keeping the bad guys out: Protecting and vaccinating deep learning with JPEG compression. May 2017.

J. Deng, W. Dong, R. Socher, L. Li, Kai Li, and Li Fei-Fei. ImageNet: A large-scale hierarchical image database. In 2009 IEEE Conference on Computer Vision and Pattern Recognition, pages 248–255, June 2009.

G. S. Dhillon, K. Azizzadenesheli, Z. C. Lipton, J. D. Bernstein, J. Kossaifi, A. Khanna, and A. Anandkumar. Stochastic activation pruning for robust adversarial defense. Feb. 2018.

J. J. DiCarlo, D. Zoccolan, and N. C. Rust. How does the brain solve visual object recognition? Neuron, 73(3):415–434, Feb. 2012.

A. Dosovitskiy, L. Beyer, A. Kolesnikov, D. Weissenborn, X. Zhai, T. Unterthiner, M. Dehghani, M. Minderer, G. Heigold, S. Gelly, J. Uszkoreit, and N. Houlsby. An image is worth 16×16 words: Transformers for image recognition at scale. Oct. 2020.

G. Elsayed, S. Shankar, B. Cheung, N. Papernot, A. Kurakin, I. Goodfellow, and J. Sohl-Dickstein. Adversarial examples that fool both computer vision and Time-Limited humans. In S. Bengio, H. Wallach, H. Larochelle, K. Grauman, N. Cesa-Bianchi, and R. Garnett, editors, Advances in Neural Information Processing Systems 31, pages 3910–3920. Curran Associates, Inc., 2018.

W. Falcon et al. Pytorch lightning. GitHub. Note: https://github.com/PyTorchLightning/pytorch-lightning, 3, 2019.

C. Federer, H. Xu, A. Fyshe, and J. Zylberberg. Improved object recognition using neural networks trained to mimic the brain’s statistical properties. Neural Netw., 2020.

F. Geiger, M. Schrimpf, T. Marques, and J. J. DiCarlo. Wiring up vision: Minimizing supervised synaptic updates needed to produce a primate ventral stream. In International Conference on Learning Representations, 2022. URL https://openreview.net/forum?id=g1SzIRLQXMM.

R. Geirhos, K. Narayanappa, B. Mitzkus, T. Thieringer, M. Bethge, F. A. Wichmann, and W. Brendel. Partial success in closing the gap between human and machine vision. Adv. Neural Inf. Process. Syst., 34, 2021.

J. Guerguiev, T. P. Lillicrap, and B. A. Richards. Towards deep learning with segregated dendrites. Elife, 6, Dec. 2017.

C. Guo, M. Rana, M. Cisse, and L. van der Maaten. Countering adversarial images using input transformations. Feb. 2018.

C. Guo, M. J. Lee, G. Leclerc, J. Dapello, Y. Rao, A. Madry, and J. J. DiCarlo. Adversarially trained neural representations may already be as robust as corresponding biological neural representations. June 2022.

H. Hasani, M. Soleymani, and H. Aghajan. Surround Modulation: A Bio-inspired Connectivity Structure for Convolutional Neural Networks. NeurIPS, (NeurIPS):15877–15888, 2019.

D. Hassabis, D. Kumaran, C. Summerfield, and M. Botvinick. Neuroscience-Inspired Artificial Intelligence. Neuron, 95(2):245–258, 2017. ISSN 10974199. doi: 10.1016/j.neuron.2017.06.011.

K. He, X. Zhang, S. Ren, and J. Sun. Delving deep into rectifiers: Surpassing human-level perfor-mance on imagenet classification. Proceedings of the IEEE International Conference on Computer Vision, 2015 Inter:1026–1034, 2015a. ISSN 15505499. doi: 10.1109/ICCV.2015.123.

K. He, X. Zhang, S. Ren, and J. Sun. Deep residual learning for image recognition. Dec. 2015b.

H. Hong, D. L. K. Yamins, N. J. Majaj, and J. J. DiCarlo. Explicit information for category-orthogonal object properties increases along the ventral stream. Nat. Neurosci., 19(4):613–622, Apr. 2016.

J. Kaplan, S. McCandlish, T. Henighan, T. B. Brown, B. Chess, R. Child, S. Gray, A. Radford, J. Wu, and D. Amodei. Scaling laws for neural language models. Jan. 2020.

K. Kar and J. J. DiCarlo. Fast recurrent processing via ventrolateral prefrontal cortex is needed by the primate ventral stream for robust core visual object recognition. Neuron, 109(1):164–176, 2021.

K. Kar, J. Kubilius, K. Schmidt, E. B. Issa, and J. J. DiCarlo. Evidence that recurrent circuits are critical to the ventral stream’s execution of core object recognition behavior. Nature neuroscience, 22(6):974–983, 2019.

S.-M. Khaligh-Razavi and N. Kriegeskorte. Deep supervised, but not unsupervised, models may explain IT cortical representation. PLoS Comput. Biol., 10(11):e1003915, Nov. 2014.

S. Kornblith, M. Norouzi, H. Lee, and G. Hinton. Similarity of neural network representations revisited. May 2019.

A. Krizhevsky, I. Sutskever, and G. E. Hinton. ImageNet classification with deep convolutional neural networks. In F. Pereira, C. J. C. Burges, L. Bottou, and K. Q. Weinberger, editors, Advances in Neural Information Processing Systems 25, pages 1097–1105. Curran Associates, Inc., 2012.

J. Kubilius, M. Schrimpf, K. Kar, H. Hong, N. J. Majaj, R. Rajalingham, E. B. Issa, P. Bashivan, J. Prescott-Roy, K. Schmidt, A. Nayebi, D. Bear, D. L. K. Yamins, and J. J. DiCarlo. Brain-Like object recognition with High-Performing shallow recurrent ANNs. Sept. 2019.

A. Kurakin, I. Goodfellow, and S. Bengio. Adversarial examples in the physical world. July 2016.

Z. Li, W. Brendel, E. Y. Walker, E. Cobos, T. Muhammad, J. Reimer, M. Bethge, F. H. Sinz, X. Pitkow, and A. S. Tolias. Learning from brains how to regularize machines. Nov. 2019.

T.-Y. Lin, M. Maire, S. Belongie, L. Bourdev, R. Girshick, J. Hays, P. Perona, D. Ramanan, C. Lawrence Zitnick, and P. Dollár. Microsoft COCO: Common objects in context. May 2014.

G. W. Lindsay and K. D. Miller. How biological attention mechanisms improve task performance in a large-scale visual system model. Elife, 7, Oct. 2018.

X. Liu, M. Cheng, H. Zhang, and C.-J. Hsieh. Towards robust neural networks via random self-ensemble. Dec. 2017.

Z. Liu, H. Mao, C.-Y. Wu, C. Feichtenhofer, T. Darrell, and S. Xie. A convnet for the 2020s. 2022.

W. Lotter, G. Kreiman, and D. Cox. Deep predictive coding networks for video prediction and unsupervised learning. May 2016.

A. Madry, A. Makelov, L. Schmidt, D. Tsipras, and A. Vladu. Towards deep learning models resistant to adversarial attacks. June 2017.

N. J. Majaj, H. Hong, E. A. Solomon, and J. J. DiCarlo. Simple learned weighted sums of inferior temporal neuronal firing rates accurately predict human core object recognition performance. J. Neurosci., 35(39):13402–13418, Sept. 2015.

A. H. Marblestone, G. Wayne, and K. P. Kording. Toward an integration of deep learning and neuroscience. Front. Comput. Neurosci., 10:94, Sept. 2016.

C. Michaelis, B. Mitzkus, R. Geirhos, E. Rusak, O. Bringmann, A. S. Ecker, M. Bethge, and W. Brendel. Benchmarking Robustness in Object Detection: Autonomous Driving when Winter is Coming. pages 1–23, 2019. URL http://arxiv.org/abs/1907.07484.

A. Nayebi and S. Ganguli. Biologically inspired protection of deep networks from adversarial attacks. Mar. 2017.

A. Nayebi, D. Bear, J. Kubilius, K. Kar, S. Ganguli, D. Sussillo, J. J. DiCarlo, and D. L. K. Yamins. Task-Driven convolutional recurrent models of the visual system. June 2018.

M.-I. Nicolae, M. Sinn, M. N. Tran, B. Buesser, A. Rawat, M. Wistuba, V. Zantedeschi, N. Baracaldo, B. Chen, H. Ludwig, I. Molloy, and B. Edwards. Adversarial robustness toolbox v1.2.0. CoRR, 1807.01069, 2018. URL https://arxiv.org/pdf/1807.01069.

A. Paszke, S. Gross, S. Chintala, G. Chanan, E. Yang, Z. DeVito, Z. Lin, A. Desmaison, L. Antiga, and A. Lerer. Automatic differentiation in PyTorch. Oct. 2017.

R. Rajalingham, K. Schmidt, and J. J. DiCarlo. Comparison of Object Recognition Behavior in Human and Monkey. Journal of Neuroscience, 35(35):12127–12136, 2015. ISSN 0270-6474. doi: 10.1523/JNEUROSCI.0573-15.2015. URL http://www.jneurosci.org/cgi/doi/10.1523/JNEUROSCI.0573-15.2015.

R. Rajalingham, E. B. Issa, P. Bashivan, K. Kar, K. Schmidt, and J. J. DiCarlo. Large-Scale, High-Resolution comparison of the core visual object recognition behavior of humans, monkeys, and State-of-the-Art deep artificial neural networks. J. Neurosci., 38(33):7255–7269, Aug. 2018.

R. Rajalingham, K. Kar, S. Sanghavi, S. Dehaene, and J. J. DiCarlo. The inferior temporal cor-tex is a potential cortical precursor of orthographic processing in untrained monkeys. Nature communications, 11(1):1–13, 2020.

A. Riedel. Bag of tricks for training brain-like deep neural networks. In Brain-Score Workshop, 2022. URL https://openreview.net/forum?id=SudzH-vWQ-c.

J. Rony, L. G. Hafemann, L. S. Oliveira, I. Ben Ayed, R. Sabourin, and E. Granger. Decoupling direction and norm for efficient Gradient-Based L2 adversarial attacks and defenses. Nov. 2018.

S. Safarani, A. Nix, K. Willeke, S. A. Cadena, K. Restivo, G. Denfield, A. S. Tolias, and F. H. Sinz. Towards robust vision by multi-task learning on monkey visual cortex. July 2021.

M. Schrimpf, J. Kubilius, H. Hong, N. J. Majaj, R. Rajalingham, E. B. Issa, K. Kar, P. Bashivan, J. Prescott-Roy, K. Schmidt, D. L. K. Yamins, and J. J. DiCarlo. Brain-Score: Which artificial neural network for object recognition is most Brain-Like? Sept. 2018.

M. Schrimpf, J. Kubilius, M. J. Lee, N. A. R. Murty, R. Ajemian, and J. J. DiCarlo. Integrative benchmarking to advance neurally mechanistic models of human intelligence. Neuron, 2020. URL https://www.cell.com/neuron/fulltext/S0896-6273(20)30605-X.

K. Simonyan and A. Zisserman. Very deep convolutional networks for Large-Scale image recognition. Sept. 2014.

F. H. Sinz, X. Pitkow, J. Reimer, M. Bethge, and A. S. Tolias. Engineering a less artificial intelligence. Neuron, 103(6):967–979, Sept. 2019.

Y. Song, T. Kim, S. Nowozin, S. Ermon, and N. Kushman. PixelDefend: Leveraging generative models to understand and defend against adversarial examples. Oct. 2017.

C. Szegedy, W. Zaremba, I. Sutskever, J. Bruna, D. Erhan, I. Goodfellow, and R. Fergus. Intriguing properties of neural networks. Dec. 2013.

C. Szegedy, W. Liu, Y. Jia, P. Sermanet, S. Reed, D. Anguelov, D. Erhan, V. Vanhoucke, and A. Rabinovich. Going deeper with convolutions. Sept. 2014.

H. Tang, M. Schrimpf, W. Lotter, C. Moerman, A. Paredes, J. Ortega Caro, W. Hardesty, D. Cox, and G. Kreiman. Recurrent computations for visual pattern completion. Proceedings of the National Academy of Sciences, 115(35):8835–8840, 2018. ISSN 0027-8424. doi: 10.1073/pnas.1719397115.

W. Xu, D. Evans, and Y. Qi. Feature squeezing: Detecting adversarial examples in deep neural networks. arXiv [cs.CV], Apr. 2017.

L. Yuan, W. Xiao, G. Dellaferrera, G. Kreiman, F. E. H. Tay, J. Feng, and M. S. Livingstone. Fooling the primate brain with minimal, targeted image manipulation. Nov. 2020.

A. M. Zador. A critique of pure learning and what artificial neural networks can learn from animal brains. Nat. Commun., 10(1):3770, Aug. 2019.

